## Supplementary Information for "Quantification of amyloid fibril polymorphism by nano-morphometry reveals the individuality of filament assembly"

Liam D. Aubrey [1†], Ben J. F. Blakeman [1†], Liisa Lutter [1], Christopher J. Serpell [2], Mick F. Tuite [1], Louise C. Serpell [3], Wei-Feng Xue [1*]

[1] Kent Fungal Group, School of Biosciences, University of Kent, CT2 7NJ, Canterbury, UK

[2] School of Physical Sciences, University of Kent, CT2 7NH, Canterbury, UK

[3] Sussex Neuroscience, School of Life Sciences, University of Sussex, BN1 6NN, Falmer, Brighton, UK

[†] Authors contributed equally to this study

Key words: amyloid / fibril structure / polymorphism / self-assembly / atomic force microscopy / image analysis

**Supplementary Table SI 1: Morphometric parameters for individual fibrils assembled from the peptides HYFNIF, RVFNIM and VIYKI.** The contour length of the section of each fibril used to estimate their morphometric parameters and the estimated radius of the cantilever tip used to image each fibril are also shown for each individual fibril. The fibril number is the index number of each of the individual fibrils and were used throughout.

| Fibril number | Length / nm | Estimated Tip-Radius / nm | Average Height, $h_{mean}$ / nm | Directional Periodic Frequency, $dpf$ / nm$^{-1}$ | Minimum Height, $h_{min}$ / nm | Maximum Height, $h_{max}$ / nm | Cross-Sectional Area, $csa$ / nm$^2$ |
|---|---|---|---|---|---|---|---|
| | | | | **HYFNIF** | | | |
| **1** | 4233.6 | 3.3 | 6.38 | -0.01133 | 5.06 | 7.97 | 30.15 |
| **2** | 2919.0 | 3.1 | 8.76 | -0.01882 | 7.78 | 9.82 | 50.60 |
| **3** | 6382.6 | 3.5 | 7.88 | -0.02459 | 7.08 | 8.62 | 44.58 |
| **4** | 6230.5 | 7.9 | 9.58 | -0.01011 | 7.79 | 11.18 | 62.38 |
| **5** | 6884.2 | 8.0 | 10.46 | -0.01351 | 9.48 | 11.32 | 82.23 |
| **6** | 1184.8 | 6.8 | 6.51 | 0.01348 | 5.74 | 7.23 | 33.06 |
| **7** | 1800.4 | 11.8 | 7.41 | -0.02497 | 6.69 | 7.94 | 42.87 |
| **8** | 863.6 | 8.7 | 6.29 | -0.01387 | 5.52 | 7.21 | 29.80 |
| **9** | 2224.9 | 7.3 | 7.42 | -0.00943 | 6.69 | 8.33 | 45.76 |
| **10** | 2222.0 | 11.2 | 9.00 | -0.01664 | 8.10 | 9.81 | 58.98 |
| **11** | 1604.3 | 12.5 | 6.39 | -0.00996 | 5.36 | 7.04 | 31.87 |
| **12** | 2353.7 | 12.7 | 6.11 | -0.01401 | 5.28 | 7.01 | 25.41 |
| **13** | 2708.0 | 9.3 | 6.94 | -0.01181 | 6.05 | 8.15 | 42.06 |
| **14** | 1785.8 | 12.1 | 7.05 | -0.00951 | 6.15 | 8.17 | 33.87 |
| **15** | 1294.0 | 9.8 | 6.96 | -0.01157 | 6.05 | 8.26 | 46.20 |
| **16** | 1056.8 | 11.4 | 6.51 | -0.01133 | 5.87 | 7.12 | 34.55 |
| **17** | 629.4 | 8.7 | 7.25 | -0.00950 | 6.42 | 8.34 | 41.34 |
| **18** | 951.5 | 12.1 | 7.50 | -0.02098 | 6.95 | 7.97 | 47.67 |
| **19** | 1074.4 | 10.5 | 7.38 | -0.00743 | 6.25 | 9.12 | 42.75 |
| **20** | 3275.9 | 12.9 | 6.56 | -0.00824 | 5.62 | 7.82 | 30.91 |
| **21** | 2470.9 | 14.0 | 6.25 | -0.00970 | 5.12 | 7.89 | 25.79 |
| **22** | 1185.7 | 12.1 | 6.41 | -0.01263 | 5.64 | 7.32 | 31.38 |
| **23** | 2775.3 | 13.3 | 6.51 | -0.01224 | 5.70 | 7.48 | 30.63 |
| **24** | 3893.7 | 12.1 | 6.78 | -0.00821 | 5.68 | 7.97 | 33.35 |
| **25** | 2227.9 | 12.0 | 6.34 | -0.01569 | 5.47 | 7.28 | 28.42 |
| **26** | 4243.1 | 11.4 | 7.45 | -0.00801 | 6.67 | 8.41 | 40.46 |
| **27** | 6256.3 | 11.7 | 7.13 | -0.00831 | 6.44 | 8.07 | 37.42 |
| **28** | 1362.6 | 12.8 | 6.89 | -0.01025 | 5.84 | 8.66 | 28.81 |
| **29** | 389.7 | 2.3 | 7.08 | -0.02556 | 6.10 | 7.94 | 31.07 |
| **30** | 799.8 | 5.8 | 5.18 | 0.00749 | 4.40 | 5.79 | 20.46 |
| **31** | 1669.9 | 2.0 | 7.63 | -0.00957 | 6.76 | 8.54 | 33.77 |
| **32** | 796.9 | 2.4 | 7.14 | -0.02380 | 6.06 | 8.00 | 33.06 |
| **33** | 896.5 | 1.9 | 7.69 | -0.01002 | 6.92 | 8.76 | 34.53 |
| **34** | 1037.1 | 1.1 | 6.04 | -0.01925 | 5.15 | 7.00 | 46.76 |
| **35** | 1869.3 | 5.6 | 5.94 | -0.01977 | 5.15 | 6.85 | 24.15 |

| | | | | | | | |
|---|---|---|---|---|---|---|---|
| **36** | 914.1 | 4.7 | 6.33 | -0.01201 | 5.54 | 7.40 | 26.40 |
| **37** | 981.5 | 4.0 | 6.05 | -0.02034 | 4.98 | 7.02 | 27.18 |
| **38** | 2088.9 | 3.3 | 6.32 | -0.01722 | 5.44 | 7.24 | 27.40 |
| **39** | 1517.6 | 5.0 | 7.11 | -0.00987 | 6.11 | 8.46 | 29.59 |
| **40** | 2504.9 | 6.7 | 7.44 | -0.01835 | 6.43 | 8.24 | 38.93 |
| **41** | 4174.8 | 7.3 | 7.72 | -0.01652 | 6.75 | 8.56 | 39.17 |
| **42** | 1391.6 | 5.6 | 6.50 | -0.01506 | 5.56 | 7.67 | 25.61 |
| **43** | 999.0 | 3.9 | 5.90 | -0.01598 | 4.98 | 6.78 | 21.07 |
| **44** | 2062.5 | 4.5 | 7.52 | -0.01598 | 6.51 | 8.62 | 35.99 |
| **45** | 2042.0 | 5.7 | 9.92 | -0.00978 | 8.06 | 12.06 | 57.88 |
| **46** | 533.2 | 7.5 | 7.38 | -0.01121 | 6.14 | 8.97 | 33.31 |
| **47** | 1880.9 | 4.1 | 7.85 | -0.01647 | 6.58 | 9.00 | 43.65 |
| **48** | 612.3 | 5.6 | 7.68 | -0.01464 | 6.15 | 9.16 | 35.66 |
| **49** | 1432.6 | 5.9 | 7.49 | -0.01115 | 6.50 | 8.64 | 34.90 |
| **50** | 3155.3 | 10.9 | 6.22 | -0.00950 | 5.40 | 7.01 | 41.08 |
| **51** | 3638.7 | 12.4 | 6.10 | -0.01373 | 5.10 | 7.16 | 44.01 |
| **52** | 5622.1 | 13.1 | 8.85 | -0.01671 | 7.84 | 9.65 | 83.03 |
| **53** | 1286.1 | 9.0 | 6.14 | -0.01474 | 5.10 | 7.10 | 31.87 |
| **54** | 726.6 | 13.0 | 6.07 | -0.01371 | 5.12 | 7.07 | 42.14 |
| **55** | 3747.1 | 12.5 | 5.44 | -0.01520 | 4.61 | 6.21 | 40.52 |
| **56** | 3295.9 | 11.5 | 8.37 | -0.01334 | 7.25 | 9.19 | 72.00 |
| **57** | 5525.4 | 11.4 | 6.23 | -0.00977 | 5.49 | 7.07 | 44.66 |
| **58** | 609.4 | 11.7 | 5.96 | -0.01471 | 5.00 | 7.01 | 40.42 |
| **59** | 1936.5 | 5.6 | 8.39 | -0.00619 | 7.41 | 10.03 | 40.26 |
| **60** | 665.1 | 8.8 | 10.38 | -0.01198 | 7.05 | 13.37 | 54.66 |
| **61** | 934.6 | 11.2 | 7.97 | -0.00854 | 6.70 | 8.95 | 46.55 |
| **62** | 1834.0 | 5.7 | 9.53 | -0.00980 | 7.02 | 12.61 | 45.83 |
| **63** | 931.8 | 4.8 | 10.22 | -0.01071 | 7.11 | 13.34 | 52.96 |
| **64** | 1711.0 | 3.8 | 7.88 | -0.01226 | 6.47 | 9.58 | 36.15 |
| **65** | 1110.4 | 3.4 | 8.73 | -0.01348 | 7.86 | 9.79 | 48.84 |
| **66** | 855.5 | 3.2 | 8.40 | -0.01283 | 7.57 | 9.22 | 46.21 |
| **67** | 3890.2 | 10.9 | 10.64 | -0.01105 | 7.40 | 13.52 | 62.20 |
| **68** | 2042.9 | 17.9 | 7.81 | -0.00636 | 6.03 | 10.78 | 37.92 |
| **69** | 2077.8 | 20.0 | 7.77 | -0.00577 | 7.05 | 8.78 | 91.02 |
| **70** | 1281.1 | 11.6 | 8.33 | -0.00857 | 7.42 | 9.73 | 50.56 |
| **71** | 2212.1 | 9.9 | 8.00 | -0.00677 | 6.18 | 11.03 | 28.84 |
| **72** | 1666.4 | 19.1 | 7.83 | -0.00659 | 7.16 | 8.83 | 95.11 |
| **73** | 1748.1 | 8.6 | 8.55 | -0.01657 | 7.79 | 9.42 | 57.09 |
| **74** | 1339.5 | 14.5 | 7.90 | -0.00820 | 5.80 | 10.53 | 28.90 |
| **75** | 1307.4 | 11.5 | 7.87 | -0.00687 | 5.93 | 10.79 | 31.94 |
| **76** | 1181.9 | 9.1 | 8.76 | -0.01013 | 7.90 | 9.60 | 59.36 |
| **77** | 2320.0 | 8.3 | 8.69 | 0.01033 | 7.80 | 9.45 | 57.57 |
| **78** | 3814.2 | 8.2 | 7.24 | -0.00996 | 5.87 | 8.86 | 33.87 |
| **79** | 1476.7 | 19.5 | 7.94 | -0.00946 | 7.20 | 8.52 | 102.43 |
| **80** | 1333.7 | 9.8 | 7.05 | -0.00823 | 5.77 | 8.73 | 33.55 |
| **81** | 1109.0 | 13.2 | 8.47 | -0.00630 | 7.22 | 10.39 | 43.35 |
| **82** | 1684.6 | 17.8 | 7.74 | 0.01542 | 6.65 | 8.67 | 77.95 |
| **83** | 2959.0 | 19.7 | 5.88 | 0.01587 | 4.86 | 6.90 | 42.55 |
| **84** | 1801.8 | 11.1 | 8.04 | 0.01608 | 7.04 | 8.91 | 58.71 |
| **85** | 6237.4 | 13.2 | 6.70 | -0.00962 | 5.83 | 7.82 | 43.78 |
| **86** | 3014.7 | 11.2 | 8.24 | -0.01525 | 6.92 | 9.06 | 51.49 |
| **87** | 2891.7 | 12.0 | 6.51 | -0.00864 | 5.74 | 7.65 | 29.87 |
| **88** | 1919.0 | 12.4 | 8.24 | -0.01770 | 7.24 | 8.91 | 54.99 |
| **89** | 1781.3 | 14.7 | 5.87 | -0.01570 | 4.98 | 6.84 | 27.54 |

| | | | | | | | |
|---|---|---|---|---|---|---|---|
| **90** | 1696.3 | 22.6 | 6.81 | -0.00883 | 5.87 | 8.17 | 51.99 |
| **91** | 1505.9 | 21.1 | 5.71 | -0.01592 | 4.79 | 6.53 | 31.82 |
| **92** | 1330.1 | 22.3 | 5.67 | -0.01426 | 4.72 | 6.46 | 32.44 |

| | | | | | | | |
|---|---|---|---|---|---|---|---|
| | | | **RVFNIM** | | | | |
| **1** | 3317.7 | 20.6 | 7.75 | -0.01295 | 6.08 | 9.25 | 53.59 |
| **2** | 2101.9 | 13.8 | 7.98 | -0.01331 | 6.15 | 9.50 | 52.72 |
| **3** | 599.2 | 13.6 | 7.65 | 0.00831 | 6.23 | 9.42 | 45.05 |
| **4** | 1026.0 | 22.0 | 7.53 | 0.01265 | 6.81 | 8.42 | 55.26 |
| **5** | 1335.8 | 23.3 | 7.48 | 0.01270 | 6.76 | 8.44 | 58.52 |
| **6** | 1034.8 | 23.6 | 7.50 | 0.01254 | 6.72 | 8.37 | 60.36 |
| **7** | 1368.0 | 21.4 | 7.51 | 0.01313 | 6.83 | 8.36 | 61.64 |
| **8** | 838.9 | 12.0 | 7.39 | -0.00171 | 4.62 | 8.17 | 53.15 |
| **9** | 780.5 | 27.9 | 5.43 | -0.00639 | 5.25 | 6.10 | 50.30 |
| **10** | 2265.4 | 20.6 | 7.53 | 0.01235 | 6.83 | 8.37 | 62.36 |
| **11** | 1961.4 | 20.6 | 7.54 | 0.01273 | 6.78 | 8.38 | 63.12 |
| **12** | 1178.0 | 14.6 | 7.68 | 0.01356 | 6.95 | 8.47 | 52.26 |
| **13** | 945.3 | 9.4 | 10.56 | -0.00845 | 8.25 | 12.43 | 77.25 |
| **14** | 2370.6 | 9.2 | 10.25 | -0.00885 | 8.10 | 12.09 | 74.65 |
| **15** | 1186.8 | 9.7 | 7.31 | 0.01430 | 6.49 | 8.23 | 42.41 |
| **16** | 2632.4 | 8.9 | 10.64 | -0.00873 | 8.33 | 12.38 | 78.72 |
| **17** | 4683.1 | 9.6 | 10.61 | -0.00832 | 8.47 | 12.51 | 73.51 |
| **18** | 619.6 | 9.8 | 7.12 | 0.00965 | 6.45 | 8.14 | 39.40 |
| **19** | 4566.4 | 10.5 | 10.73 | -0.00860 | 8.38 | 12.82 | 84.25 |
| **20** | 802.6 | 10.1 | 7.17 | 0.00995 | 6.46 | 8.06 | 47.28 |
| **21** | 3126.0 | 9.9 | 7.30 | 0.01311 | 6.63 | 8.15 | 38.89 |
| **22** | 1813.5 | 8.0 | 10.47 | -0.00826 | 8.37 | 12.19 | 70.84 |
| **23** | 3011.8 | 8.2 | 10.69 | -0.00863 | 8.57 | 12.58 | 74.25 |
| **24** | 1573.3 | 9.5 | 10.67 | -0.00826 | 8.32 | 12.51 | 75.44 |
| **25** | 1579.1 | 10.7 | 10.76 | -0.00886 | 8.38 | 12.49 | 83.57 |
| **26** | 3791.1 | 8.5 | 10.75 | -0.00844 | 8.46 | 12.76 | 78.81 |
| **27** | 1652.4 | 7.3 | 10.31 | -0.00786 | 7.94 | 12.54 | 64.46 |
| **28** | 2179.7 | 7.1 | 10.54 | -0.00917 | 8.47 | 12.69 | 69.99 |
| **29** | 1174.8 | 7.1 | 10.73 | -0.00850 | 8.53 | 12.74 | 72.98 |
| **30** | 952.2 | 8.6 | 9.54 | 0.00036 | 7.45 | 11.69 | 47.97 |
| **31** | 1210.0 | 16.4 | 11.02 | -0.00990 | 8.51 | 13.49 | 82.12 |
| **32** | 3410.2 | 6.0 | 10.24 | 0.00703 | 9.25 | 11.47 | 68.18 |
| **33** | 1113.3 | 6.0 | 8.33 | 0.00628 | 7.25 | 9.57 | 45.06 |
| **34** | 1078.1 | 6.4 | 8.48 | 0.01018 | 7.36 | 9.84 | 40.70 |
| **35** | 3020.5 | 8.3 | 9.53 | 0.00629 | 7.80 | 11.41 | 52.48 |
| **36** | 1388.7 | 8.6 | 10.45 | 0.00862 | 9.22 | 11.62 | 73.12 |
| **37** | 1324.3 | 12.0 | 9.76 | 0.00452 | 7.17 | 14.43 | 46.67 |
| **38** | 1505.9 | 16.7 | 10.67 | -0.00929 | 8.15 | 12.92 | 77.57 |
| **39** | 3087.2 | 8.1 | 11.55 | -0.00777 | 9.30 | 13.68 | 86.29 |
| **40** | 1765.3 | 6.8 | 11.58 | -0.00792 | 9.10 | 13.68 | 86.69 |
| **41** | 922.3 | 11.6 | 11.39 | -0.00866 | 9.10 | 13.32 | 92.91 |
| **42** | 4920.1 | 10.3 | 11.36 | -0.00792 | 9.17 | 13.38 | 86.04 |
| **43** | 1809.3 | 15.9 | 7.88 | -0.01160 | 6.19 | 9.72 | 40.10 |
| **44** | 4380.3 | 13.8 | 6.80 | 0.00890 | 6.06 | 7.36 | 43.37 |
| **45** | 3154.7 | 13.7 | 8.24 | -0.01109 | 6.54 | 10.06 | 38.83 |
| **46** | 1537.9 | 12.4 | 7.83 | -0.01234 | 6.10 | 9.50 | 38.63 |
| **47** | 2909.5 | 10.7 | 7.19 | -0.01305 | 5.68 | 9.12 | 33.28 |
| **48** | 4561.5 | 5.1 | 8.51 | 0.01030 | 7.63 | 9.50 | 47.76 |
| **49** | 4450.2 | 8.1 | 7.28 | -0.04380 | 6.68 | 7.83 | 41.83 |

| | | | | | | | |
|---|---|---|---|---|---|---|---|
| **50** | 6172.9 | 3.5 | 8.53 | 0.00939 | 6.75 | 10.79 | 40.14 |
| **51** | 6750.0 | 3.5 | 12.14 | -0.00933 | 10.43 | 13.98 | 86.67 |
| **52** | 668.0 | 3.7 | 10.28 | 0.00895 | 9.47 | 11.50 | 60.32 |
| **53** | 5168.0 | 4.1 | 10.08 | 0.01006 | 9.23 | 11.23 | 62.88 |
| **54** | 3966.8 | 4.3 | 10.17 | 0.00857 | 9.28 | 11.35 | 59.71 |
| **55** | 1016.6 | 4.4 | 10.09 | 0.00982 | 9.18 | 11.20 | 67.10 |
| **56** | 3366.3 | 7.4 | 10.38 | 0.00624 | 7.38 | 15.34 | 43.17 |
| **57** | 2437.5 | 4.0 | 10.01 | 0.00984 | 9.19 | 11.18 | 60.20 |
| **58** | 3799.8 | 4.2 | 7.78 | 0.00605 | 7.33 | 8.55 | 42.93 |
| **59** | 10898.5 | 2.4 | 10.01 | 0.01000 | 9.17 | 11.14 | 65.11 |
| **60** | 1628.9 | 4.2 | 9.98 | 0.01104 | 9.17 | 11.21 | 59.07 |
| **61** | 738.3 | 2.6 | 7.81 | -0.01079 | 7.48 | 8.55 | 45.57 |
| **62** | 963.9 | 2.5 | 7.83 | 0.00621 | 7.39 | 8.52 | 45.64 |
| **63** | 3293.0 | 2.5 | 7.94 | -0.01730 | 7.39 | 8.56 | 47.36 |
| **64** | 1292.0 | 4.3 | 9.98 | 0.01081 | 9.18 | 11.14 | 60.61 |
| **65** | 1895.5 | 3.6 | 8.91 | -0.00896 | 7.75 | 10.18 | 49.54 |
| **66** | 1558.6 | 4.9 | 10.11 | 0.00897 | 9.19 | 11.19 | 66.45 |
| **67** | 6820.3 | 4.5 | 9.87 | 0.00718 | 8.99 | 10.96 | 62.40 |
| **68** | 5302.2 | 6.0 | 9.86 | 0.00324 | 2.77 | 10.65 | 74.72 |
| **69** | 1199.3 | 5.5 | 6.62 | -0.02746 | 5.88 | 7.32 | 32.52 |
| **70** | 1733.4 | 7.8 | 9.14 | -0.00865 | 7.14 | 11.39 | 52.15 |
| **71** | 1277.4 | 5.6 | 8.16 | 0.00703 | 6.50 | 10.53 | 33.97 |
| **72** | 5956.1 | 4.3 | 10.18 | 0.01007 | 9.21 | 11.38 | 75.36 |
| **73** | 530.3 | 15.2 | 3.34 | -0.01315 | 2.83 | 4.15 | 8.11 |
| **74** | 865.8 | 7.3 | 4.33 | 0.01614 | 3.91 | 5.12 | 12.65 |
| **75** | 5804.1 | 6.8 | 9.17 | 0.00964 | 7.86 | 10.65 | 51.21 |
| **76** | 3022.0 | 9.4 | 5.74 | -0.00331 | 5.05 | 6.54 | 19.74 |
| **77** | 797.7 | 8.7 | 6.67 | -0.02747 | 5.86 | 7.42 | 32.31 |
| **78** | 6887.7 | 7.2 | 10.92 | -0.00145 | 9.55 | 12.82 | 85.12 |
| **79** | 3433.6 | 11.8 | 10.41 | -0.01847 | 5.91 | 13.51 | 64.64 |
| **80** | 16708.1 | 15.3 | 9.84 | 0.00934 | 8.89 | 10.74 | 100.25 |
| **81** | 1790.1 | 18.3 | 8.86 | 0.01005 | 7.90 | 9.74 | 95.32 |
| **82** | 7555.8 | 17.8 | 9.40 | -0.00939 | 8.38 | 10.64 | 95.63 |
| **83** | 2493.2 | 9.8 | 7.46 | 0.00801 | 5.51 | 10.43 | 42.50 |
| **84** | 5311.6 | 7.9 | 11.66 | 0.00903 | 9.90 | 13.50 | 93.44 |
| **85** | 2159.2 | 8.1 | 7.30 | 0.00879 | 5.62 | 9.85 | 31.94 |
| **86** | 4060.6 | 10.8 | 8.56 | -0.00763 | 6.96 | 10.59 | 51.36 |
| **87** | 1928.1 | 12.9 | 7.27 | 0.00311 | 6.15 | 9.40 | 31.21 |
| **88** | 949.2 | 13.6 | 8.32 | -0.01367 | 6.99 | 9.84 | 43.15 |
| **89** | 6758.9 | 11.1 | 7.24 | -0.01257 | 6.06 | 8.72 | 39.55 |
| | | | | **VIYKI** | | | |
| **1** | 1306.7 | 12.8 | 9.56 | -0.00687 | 8.61 | 10.77 | 75.36 |
| **2** | 3902.4 | 14.1 | 9.63 | -0.00666 | 8.80 | 10.61 | 83.29 |
| **3** | 2768.6 | 8.5 | 9.82 | -0.00650 | 9.04 | 10.60 | 72.35 |
| **4** | 3770.5 | 6.1 | 9.76 | -0.02147 | 8.99 | 10.48 | 59.62 |
| **5** | 2941.4 | 13.3 | 9.45 | -0.02581 | 8.67 | 10.18 | 97.81 |
| **6** | 5411.1 | 12.2 | 9.64 | -0.01515 | 8.86 | 10.38 | 83.65 |
| **7** | 3014.7 | 15.3 | 9.91 | 0.01890 | 9.26 | 11.03 | 84.79 |
| **8** | 3969.8 | 4.8 | 10.49 | -0.02241 | 9.82 | 11.15 | 86.68 |
| **9** | 1869.2 | 11.9 | 11.25 | 0.01443 | 10.37 | 12.40 | 105.72 |
| **10** | 1460.5 | 9.9 | 9.54 | -0.02392 | 8.86 | 10.14 | 83.66 |
| **11** | 3952.3 | 7.7 | 9.67 | -0.01416 | 8.93 | 10.29 | 72.27 |
| **12** | 1983.5 | 9.1 | 9.60 | -0.00806 | 8.76 | 10.36 | 70.86 |

| | | | | | | | |
|---|---|---|---|---|---|---|---|
| **13** | 6403.2 | 11.3 | 9.48 | -0.01561 | 8.72 | 10.16 | 84.75 |
| **14** | 872.8 | 8.8 | 9.62 | -0.02173 | 8.80 | 10.25 | 71.63 |
| **15** | 2708.9 | 8.9 | 10.50 | -0.01365 | 9.69 | 11.72 | 66.02 |
| **16** | 814.1 | 5.4 | 9.69 | -0.01348 | 9.02 | 10.38 | 55.49 |
| **17** | 5558.3 | 2.5 | 9.69 | -0.01367 | 9.03 | 10.33 | 64.84 |
| **18** | 951.8 | 4.8 | 10.22 | -0.02307 | 9.62 | 10.87 | 70.11 |
| **19** | 6946.5 | 2.2 | 9.46 | -0.00648 | 7.66 | 10.76 | 57.19 |
| **20** | 3951.3 | 6.6 | 9.49 | -0.01012 | 7.82 | 11.09 | 59.94 |
| **21** | 4168.0 | 7.7 | 10.03 | -0.02134 | 9.29 | 10.68 | 69.67 |
| **22** | 4050.8 | 8.5 | 11.01 | -0.00642 | 9.76 | 12.43 | 77.84 |
| **23** | 2574.6 | 7.5 | 10.38 | -0.01513 | 9.57 | 11.10 | 76.78 |
| **24** | 1801.3 | 7.6 | 10.31 | -0.01497 | 9.61 | 11.00 | 75.82 |
| **25** | 3488.4 | 10.2 | 9.80 | -0.01261 | 9.08 | 10.51 | 65.85 |
| **26** | 2562.4 | 8.5 | 9.87 | -0.00546 | 8.01 | 11.01 | 60.63 |
| **27** | 1144.3 | 10.8 | 9.74 | -0.00523 | 8.66 | 11.30 | 53.32 |
| **28** | 3482.5 | 10.3 | 9.87 | -0.01808 | 9.16 | 10.53 | 67.28 |
| **29** | 704.7 | 11.7 | 10.02 | -0.01131 | 9.07 | 11.40 | 58.51 |
| **30** | 3307.8 | 9.5 | 9.88 | -0.01058 | 9.16 | 10.68 | 68.44 |
| **31** | 2402.2 | 8.5 | 9.97 | -0.00499 | 8.13 | 11.25 | 58.61 |
| **32** | 5220.7 | 5.3 | 14.82 | -0.00996 | 13.65 | 16.08 | 155.04 |
| **33** | 5677.8 | 5.4 | 9.46 | -0.00951 | 7.19 | 11.27 | 128.53 |
| **34** | 12767.6 | 5.3 | 14.13 | -0.00070 | 12.86 | 16.25 | 123.08 |
| **35** | 1895.5 | 5.3 | 8.30 | -0.01212 | 6.54 | 9.95 | 52.77 |
| **36** | 4488.3 | 5.9 | 12.70 | -0.01158 | 11.09 | 14.07 | 114.90 |
| **37** | 3580.1 | 5.2 | 14.05 | -0.00503 | 13.08 | 15.21 | 142.31 |
| **38** | 3240.2 | 3.0 | 12.99 | -0.01018 | 11.64 | 14.18 | 118.56 |
| **39** | 2050.8 | 2.9 | 12.90 | -0.01023 | 11.71 | 13.97 | 118.45 |
| **40** | 1957.0 | 4.7 | 9.12 | -0.00766 | 7.60 | 10.75 | 59.94 |
| **41** | 1716.8 | 5.8 | 14.19 | -0.01571 | 13.44 | 14.92 | 149.40 |
| **42** | 5619.2 | 9.0 | 14.10 | -0.02081 | 12.58 | 15.80 | 172.76 |
| **43** | 1804.8 | 4.8 | 12.82 | -0.01162 | 11.76 | 13.53 | 121.96 |
| **44** | 3958.0 | 9.6 | 14.02 | -0.00934 | 13.29 | 14.81 | 153.36 |
| **45** | 2836.0 | 9.0 | 12.62 | -0.01022 | 10.16 | 14.29 | 149.29 |
| **46** | 665.0 | 5.0 | 12.44 | -0.00899 | 9.21 | 14.17 | 120.73 |
| **47** | 1198.3 | 9.3 | 12.91 | -0.00999 | 11.88 | 13.78 | 126.90 |
| **48** | 1207.0 | 3.1 | 8.47 | -0.01488 | 6.75 | 9.97 | 44.71 |
| **49** | 770.5 | 5.7 | 8.19 | -0.01425 | 6.64 | 9.71 | 51.21 |
| **50** | 4133.8 | 5.7 | 8.68 | -0.01427 | 7.09 | 10.04 | 54.75 |
| **51** | 2097.7 | 5.7 | 8.48 | -0.01238 | 6.89 | 10.05 | 54.08 |
| **52** | 565.4 | 7.8 | 6.08 | -0.01585 | 5.36 | 7.05 | 56.46 |
| **53** | 4749.0 | 3.5 | 6.23 | 0.00168 | 5.54 | 7.13 | 30.06 |
| **54** | 1271.5 | 2.6 | 8.55 | -0.01491 | 7.08 | 9.90 | 47.08 |
| **55** | 823.3 | 2.9 | 8.53 | -0.01334 | 6.94 | 9.98 | 45.39 |
| **56** | 1511.7 | 1.1 | 6.27 | -0.03569 | 5.71 | 6.87 | 48.89 |
| **57** | 477.5 | 3.3 | 8.47 | -0.01460 | 7.02 | 9.45 | 42.41 |
| **58** | 10878.0 | 12.8 | 8.80 | -0.01562 | 7.49 | 9.99 | 64.85 |
| **59** | 2214.9 | 10.6 | 8.94 | -0.01443 | 7.59 | 10.36 | 59.29 |
| **60** | 3012.7 | 12.2 | 9.48 | -0.01228 | 7.80 | 11.06 | 61.31 |
| **61** | 1317.5 | 11.5 | 6.73 | -0.01591 | 6.20 | 7.27 | 35.32 |
| **62** | 1461.0 | 9.2 | 8.38 | -0.01230 | 6.84 | 10.05 | 55.94 |
| **63** | 6555.3 | 12.1 | 6.68 | -0.03752 | 6.22 | 7.37 | 37.12 |
| **64** | 3340.6 | 8.6 | 9.34 | -0.01017 | 7.58 | 10.90 | 64.45 |
| **65** | 5198.4 | 3.4 | 8.03 | -0.01346 | 6.63 | 9.38 | 44.62 |
| **66** | 1679.1 | 5.2 | 8.39 | -0.01249 | 6.96 | 9.96 | 44.54 |

| | | | | | | | |
|---|---|---|---|---|---|---|---|
| **67** | 2332.5 | 2.8 | 7.31 | -0.01413 | 6.72 | 8.33 | 38.99 |
| **68** | 2124.5 | 4.9 | 9.04 | -0.01411 | 7.53 | 10.67 | 56.58 |
| **69** | 7116.3 | 16.1 | 11.82 | -0.01222 | 10.97 | 12.74 | 111.69 |
| **70** | 2097.7 | 16.7 | 10.19 | -0.00714 | 9.14 | 11.36 | 86.84 |
| **71** | 1333.0 | 4.3 | 6.46 | -0.03744 | 5.88 | 7.18 | 31.12 |
| **72** | 2097.7 | 4.4 | 6.94 | -0.01762 | 6.35 | 7.99 | 35.76 |
| **73** | 1048.9 | 4.5 | 6.42 | 0.03711 | 5.95 | 7.13 | 48.25 |
| **74** | 1145.5 | 4.6 | 6.51 | -0.03834 | 5.96 | 7.14 | 32.16 |
| **75** | 2774.4 | 4.6 | 6.46 | -0.03781 | 5.96 | 7.08 | 29.69 |
| **76** | 5845.3 | 11.1 | 6.16 | -0.01778 | 5.36 | 7.13 | 35.51 |
| **77** | 1549.6 | 7.2 | 6.24 | 0.03739 | 5.74 | 6.86 | 36.13 |
| **78** | 846.7 | 5.3 | 8.81 | -0.01532 | 7.47 | 10.10 | 51.79 |
| **79** | 1974.6 | 6.6 | 6.52 | -0.03693 | 5.97 | 7.29 | 35.61 |
| **80** | 6820.4 | 4.2 | 7.11 | -0.01583 | 6.43 | 8.04 | 37.61 |
| **81** | 1541.0 | 5.1 | 9.04 | -0.01556 | 7.76 | 10.20 | 60.05 |
| **82** | 2862.3 | 5.0 | 6.61 | -0.01431 | 6.04 | 7.34 | 30.01 |
| **83** | 3518.6 | 5.8 | 8.20 | -0.01449 | 6.70 | 9.66 | 45.13 |
| **84** | 2493.2 | 5.0 | 6.50 | -0.03646 | 5.95 | 7.11 | 29.02 |
| **85** | 1095.7 | 5.5 | 8.30 | -0.01457 | 6.84 | 9.69 | 47.00 |

#### HYFNIF

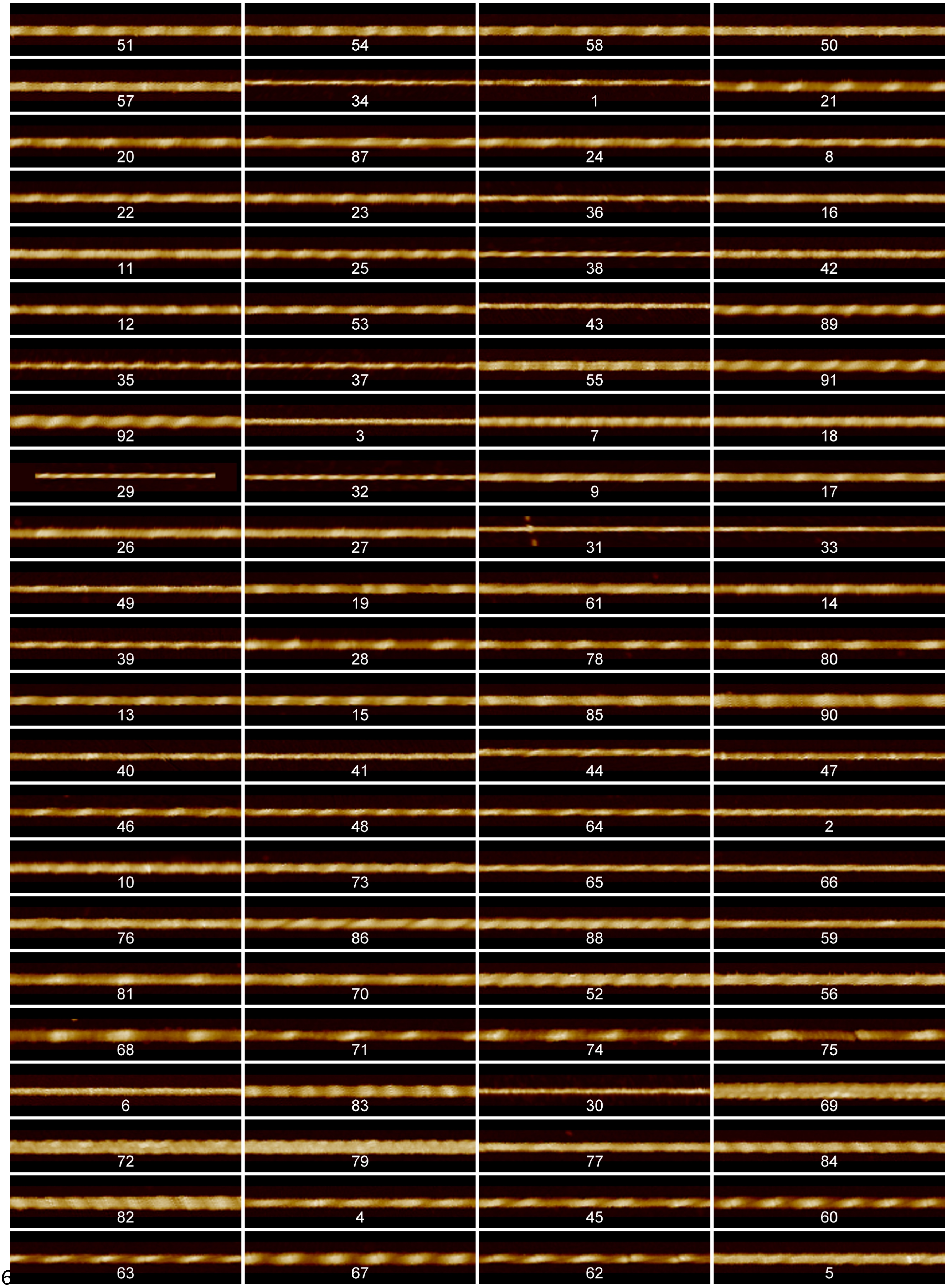
51
54
58
50
57
34
1
21
20
87
24
8
22
23
36
16
11
25
38
42
12
53
43
89
35
37
55
91
92
3
7
18
29
32
9
17
26
27
31
33
49
19
61
14
39
28
78
80
13
15
85
90
40
41
44
47
46
48
64
2
10
73
65
66
76
86
88
59
81
70
52
56
68
71
74
75
6
83
30
69
72
79
77
84
82
4
45
60
63
67
62
5

#### RVFNIM

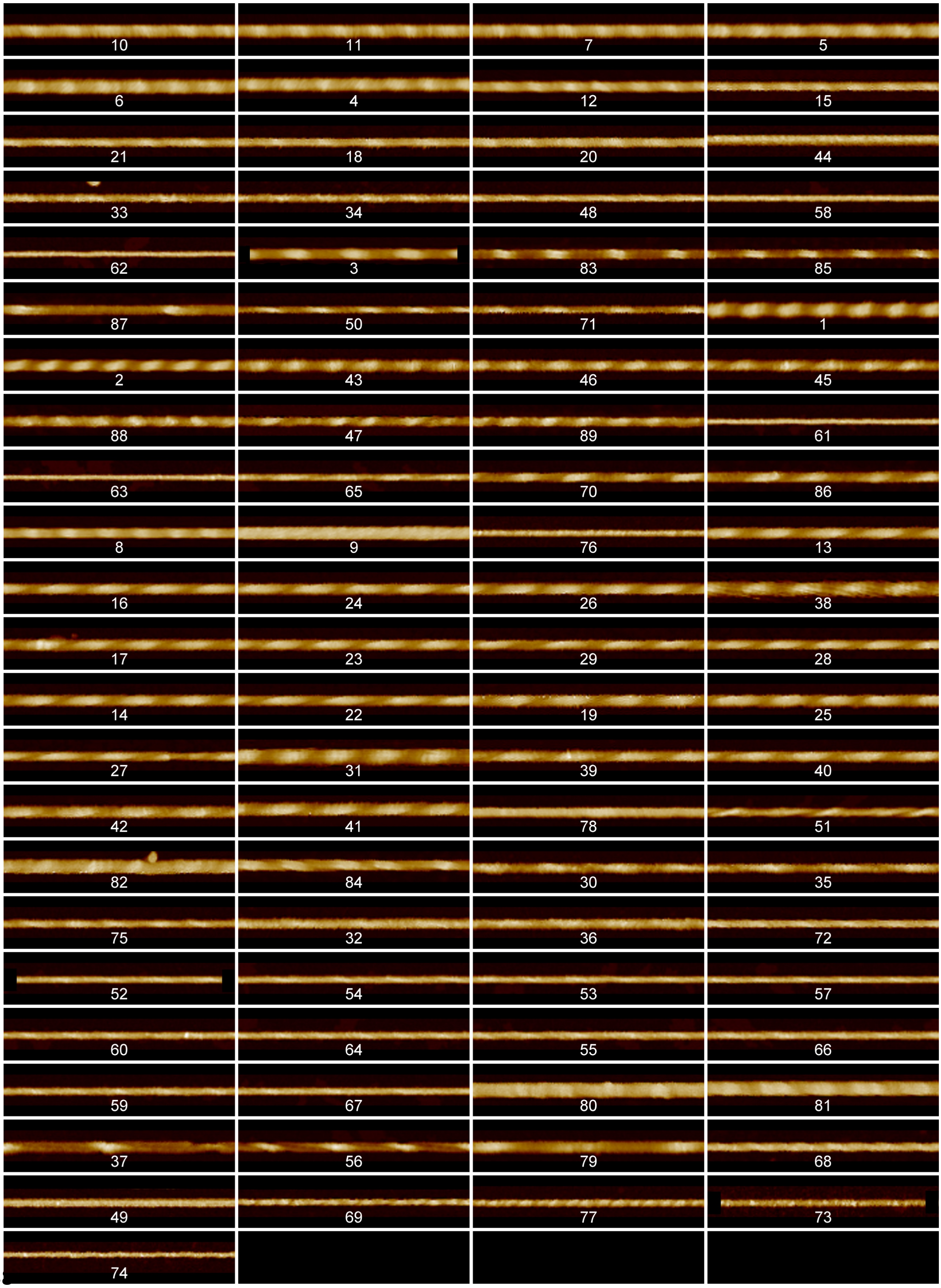
10
11
7
5
6
4
12
15
21
18
20
44
33
34
48
58
62
3
83
85
87
50
71
1
2
43
46
45
88
47
89
61
63
65
70
86
8
9
76
13
16
24
26
38
17
23
29
28
14
22
19
25
27
31
39
40
42
41
78
51
82
84
30
35
75
32
36
72
52
54
53
57
60
64
55
66
59
67
80
81
37
56
79
68
49
69
77
73
74

#### VIYKI

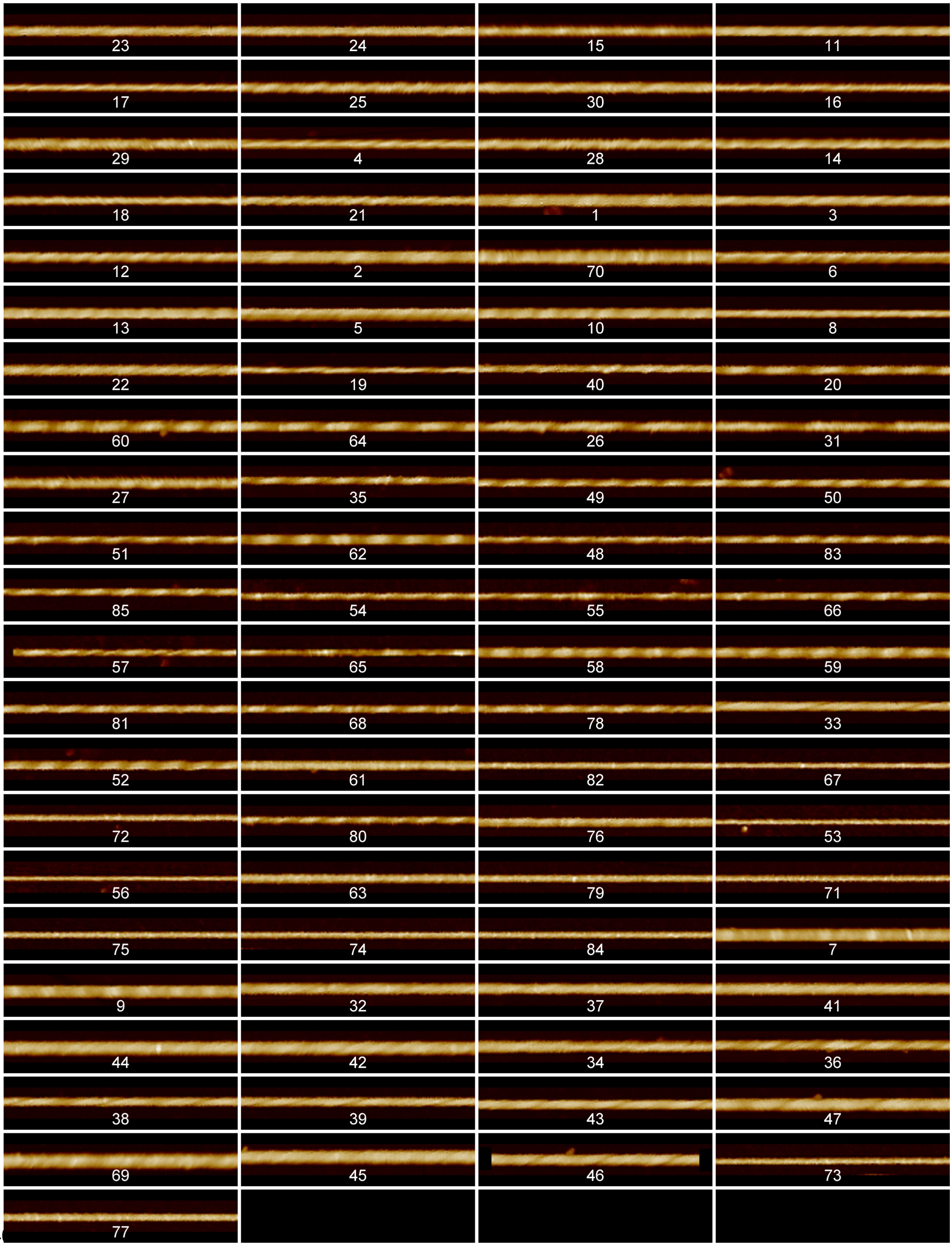
23
24
15
11
17
25
30
16
29
4
28
14
18
21
1
3
12
2
70
6
13
5
10
8
22
19
40
20
60
64
26
31
27
35
49
50
51
62
48
83
85
54
55
66
57
65
58
59
81
68
78
33
52
61
82
67
72
80
76
53
56
63
79
71
75
74
84
7
9
32
37
41
44
42
34
36
38
39
43
47
69
45
46
73
77

**Supplementary Figure SI 1: AFM image data sets for all of the individually traced fibrils.** All fibrils displayed here were straightened and they were cropped to 500 nm segments for visualisation here if the contour length is longer than 500 nm. No further processing occurred. The fibrils are shown with their individual index numbers (see Supplementary Table SI 1), and are arranged by similarity (see Figure 6 and Supplementary Figure 6).

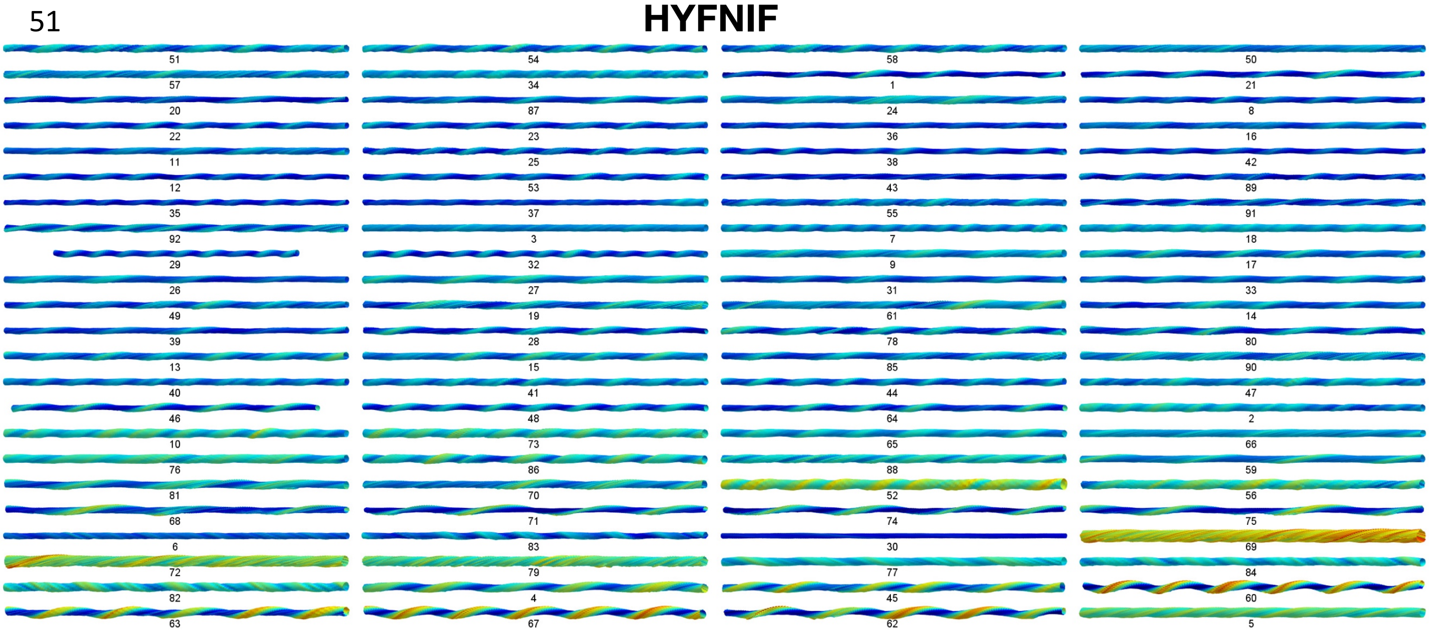
HYFNIF

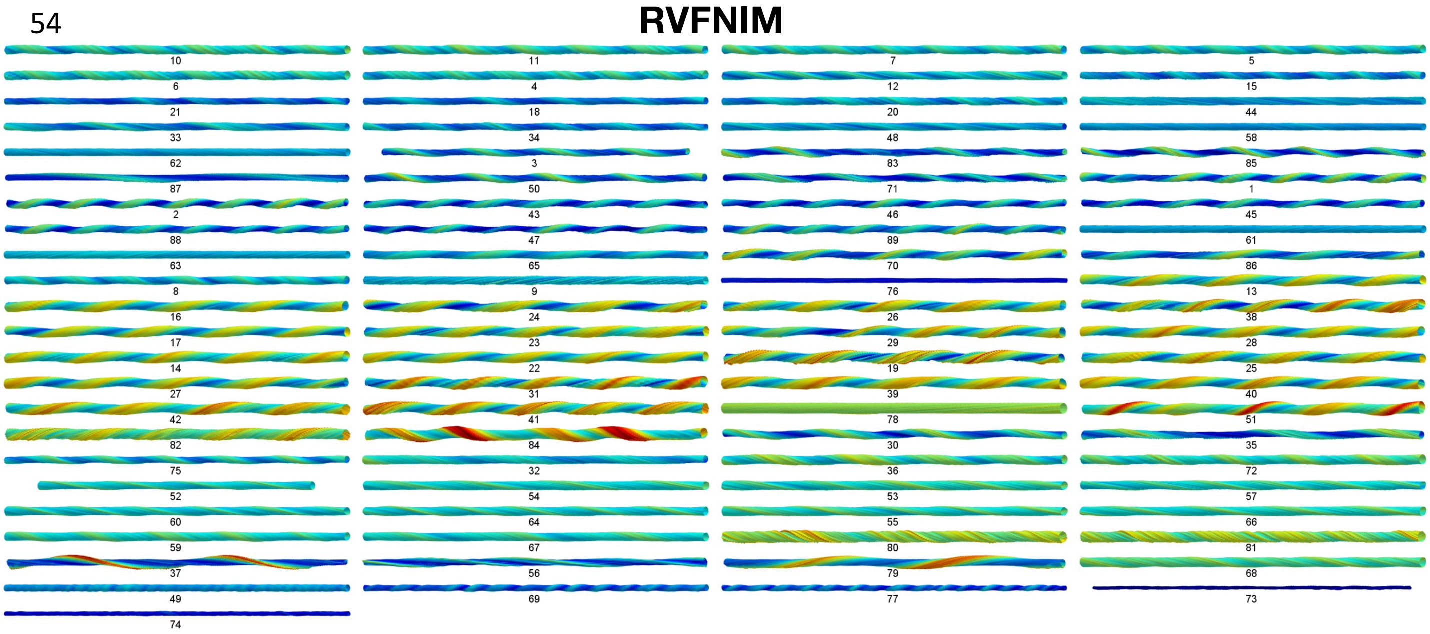
RVFNIM

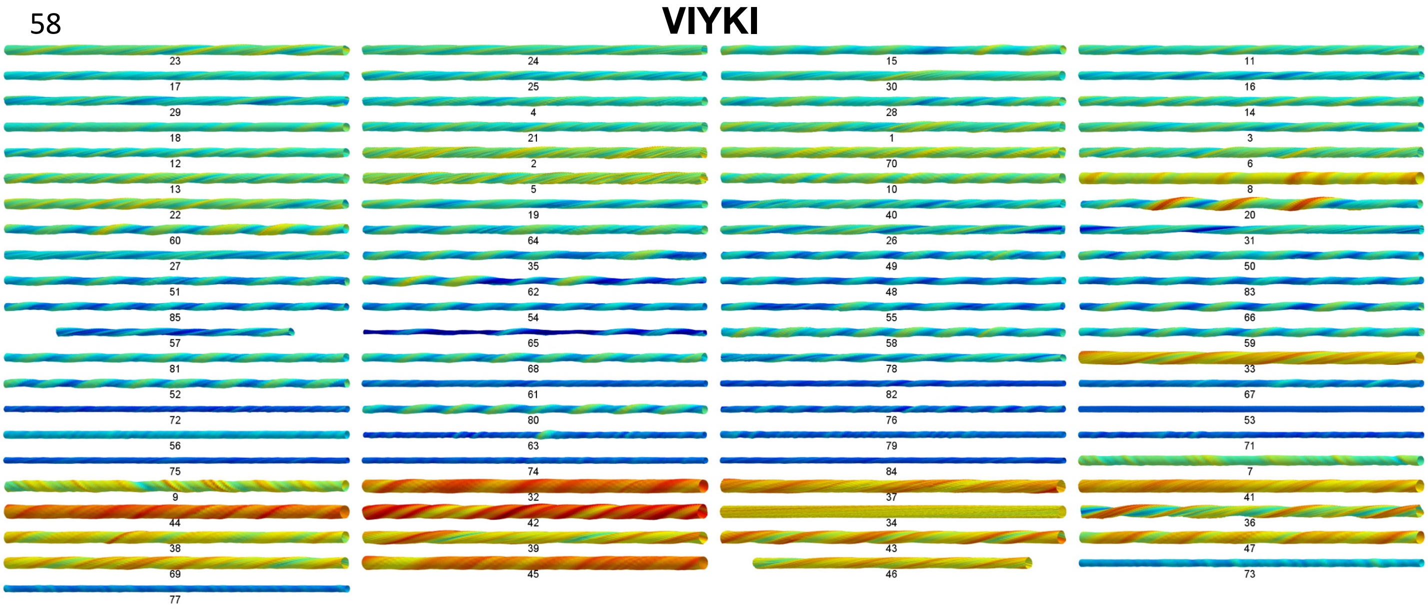

**Supplementary Figure SI 2: All 3D models of the Waltz peptide assemblies reconstructed from the AFM data sets.** Each fibril in the three data sets was reconstructed as a 3D model using information and coordinates extracted directly from the AFM data. All of the models are shown with identical scale. All fibril models displayed here are cropped to 500 nm segments for visualisation here if the contour length is longer than 500 nm. The models are shown with their individual index numbers used throughout (see Supplementary Table SI 1), are arranged by similarity (see Figure 6 and Supplementary Figure 6), and are placed in the same order as Supplementary Table SI 2.

### HYFNIF

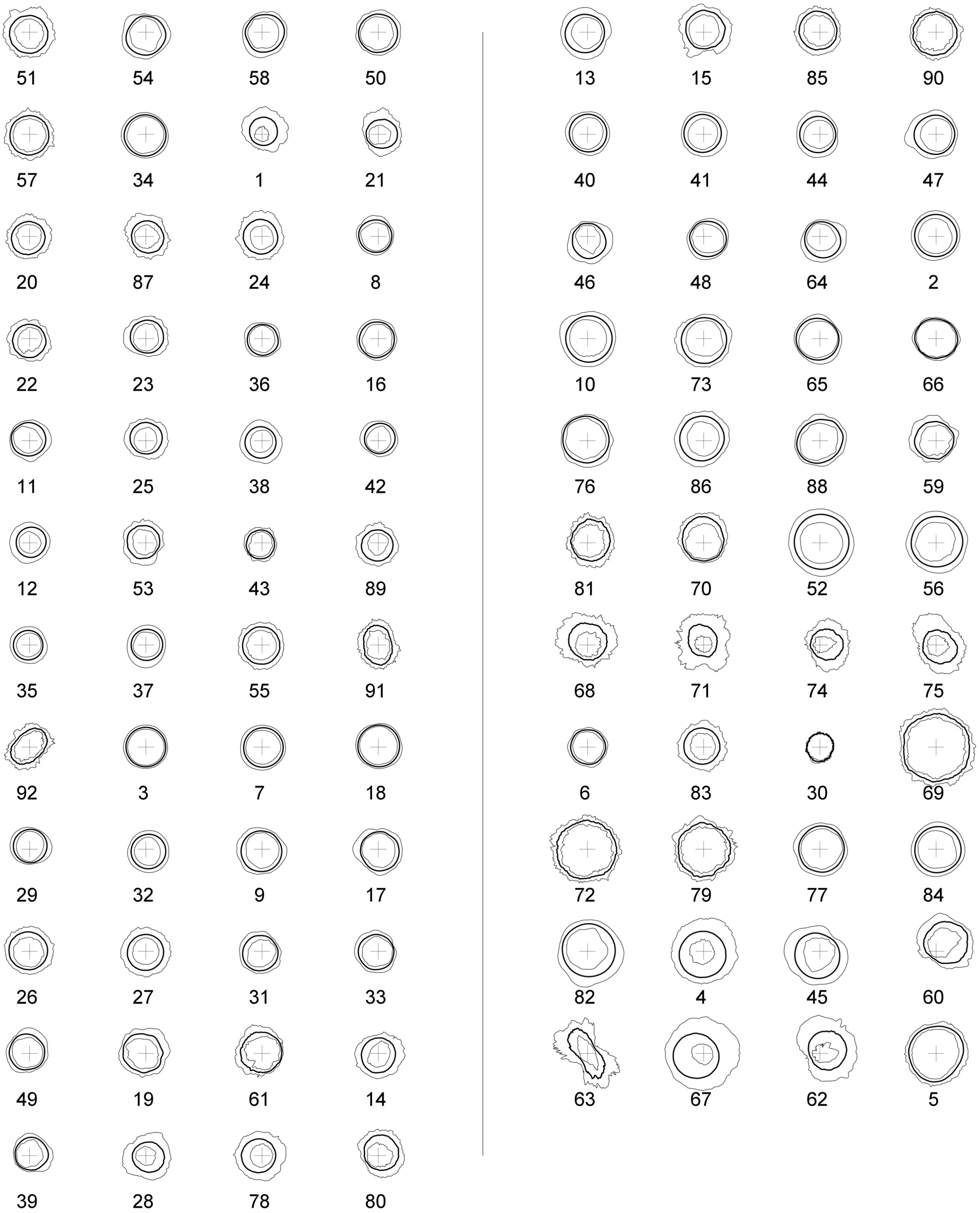
51 54 58 50
57 34 1 21
20 87 24 8
22 23 36 16
11 25 38 42
12 53 43 89
35 37 55 91
92 3 7 18
29 32 9 17
26 27 31 33
49 19 61 14
39 28 78 80
13 15 85 90
40 41 44 47
46 48 64 2
10 73 65 66
76 86 88 59
81 70 52 56
68 71 74 75
6 83 30 69
72 79 77 84
82 4 45 60
63 67 62 5

#### RVFNIM

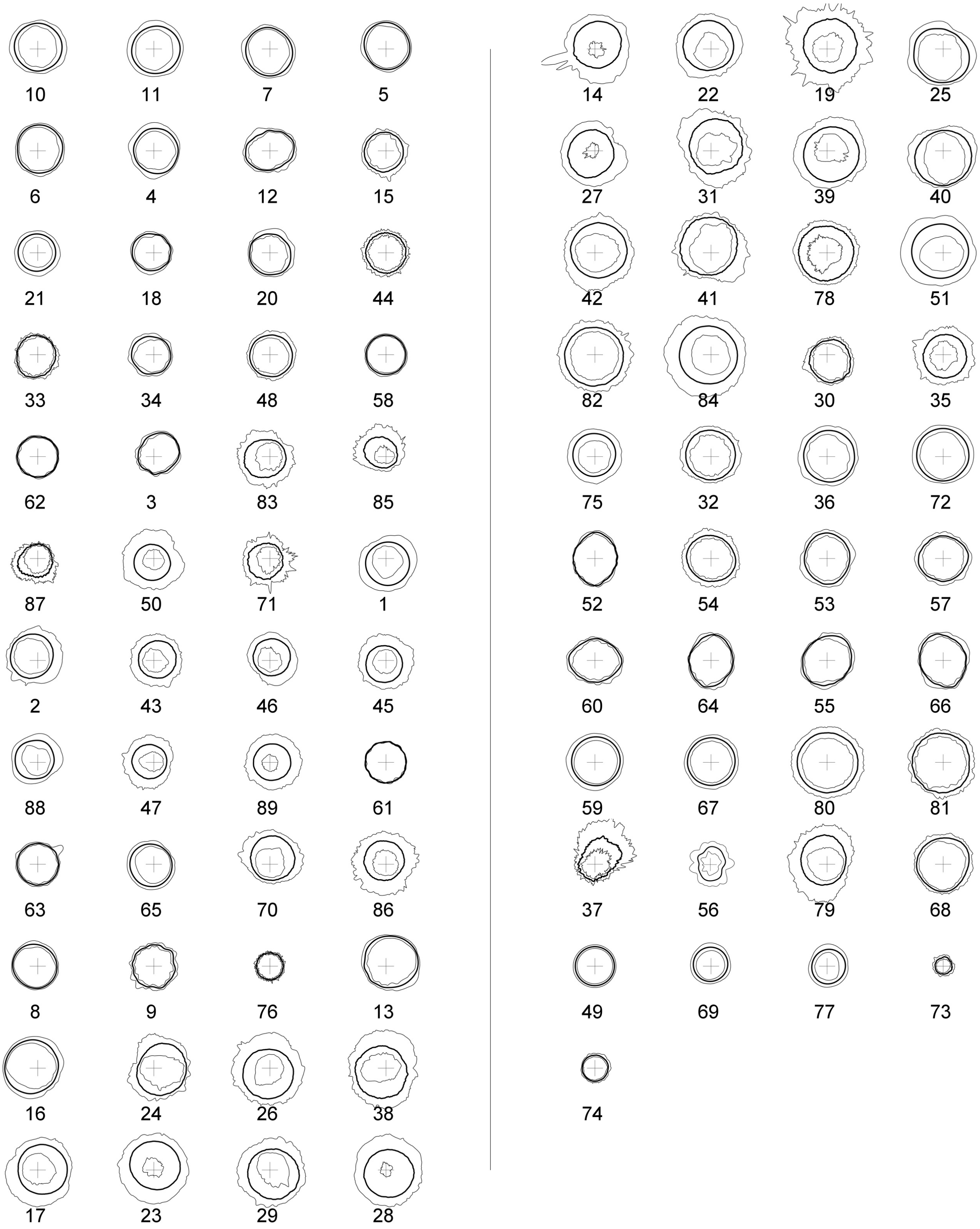
10 11 7 5 14 22 19 25
6 4 12 15 27 31 39 40
21 18 20 44 42 41 78 51
33 34 48 58 82 84 30 35
62 3 83 85 75 32 36 72
87 50 71 1 52 54 53 57
2 43 46 45 60 64 55 66
88 47 89 61 59 67 80 81
63 65 70 86 37 56 79 68
8 9 76 13 49 69 77 73
16 24 26 38 74
17 23 29 28

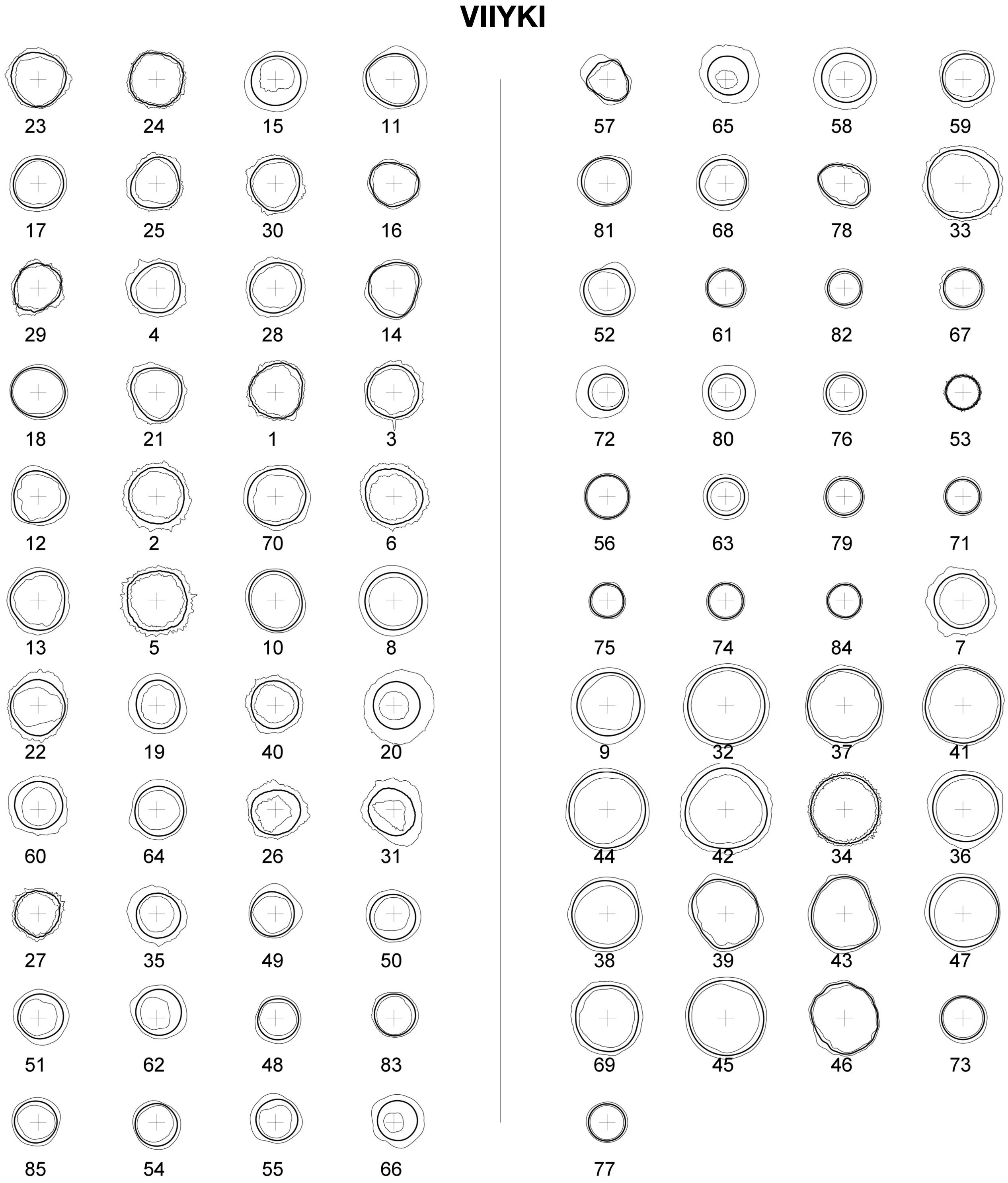

**Supplementary Figure SI 3: Average cross-sectional areas of all 3D models of the Waltz peptide assemblies reconstructed from the AFM data sets.** For each fibril 3D model shown in Supplementary Figure SI 3, the average cross-section for that individual fibril is shown as thick solid line. The thin solid lines inside and outside the average cross-sections represent the minimum and the maximum cross-sections, respectively. They reflect the variations observed in the fibril cross-sections. All of the models are shown with identical scale. The centre cross in each of the model cross-section represent their screw axis, and the length of the horizontal and vertical lines of the cross represent the length of 4 nm. The model cross-sections are shown with their individual index numbers used throughout (see Supplementary Table SI 1), are arranged by similarity (see Figure 6 and Supplementary Figure 6), and placed in the same order as Supplementary Table SI 2.

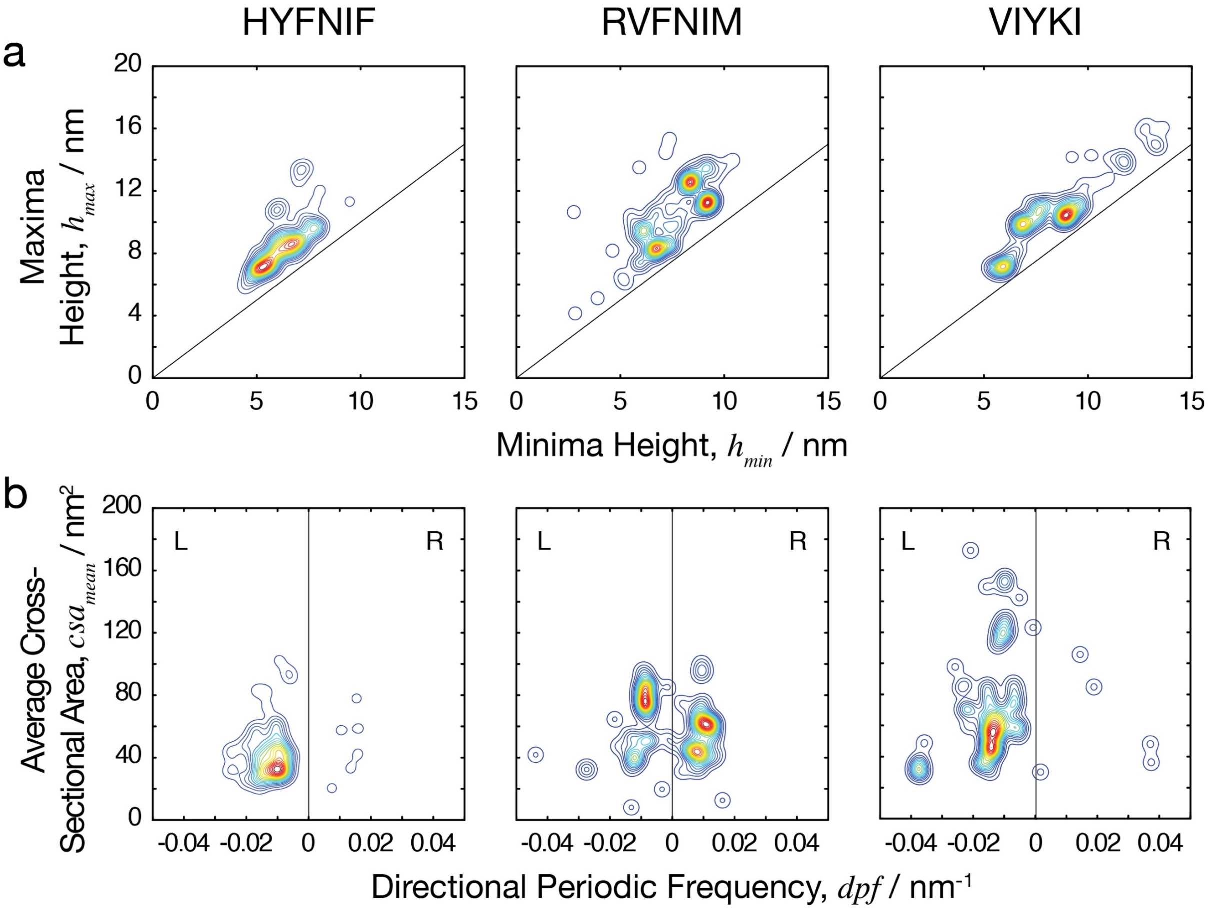

**Supplementary Figure SI 4: The heterogeneity of the Waltz peptide assemblies.** (a) The maximum height of the fibrils plotted against the minimum height of the fibrils. (b) The average cross-sectional area of the fibrils plotted against the number of repeating units per nm, with negative and positive values to distinguish handedness (directional periodic frequency, *dpf*). The data is represented as a 2D histogram and visualised as a contour map, where the colouring represents the density of the data-points.

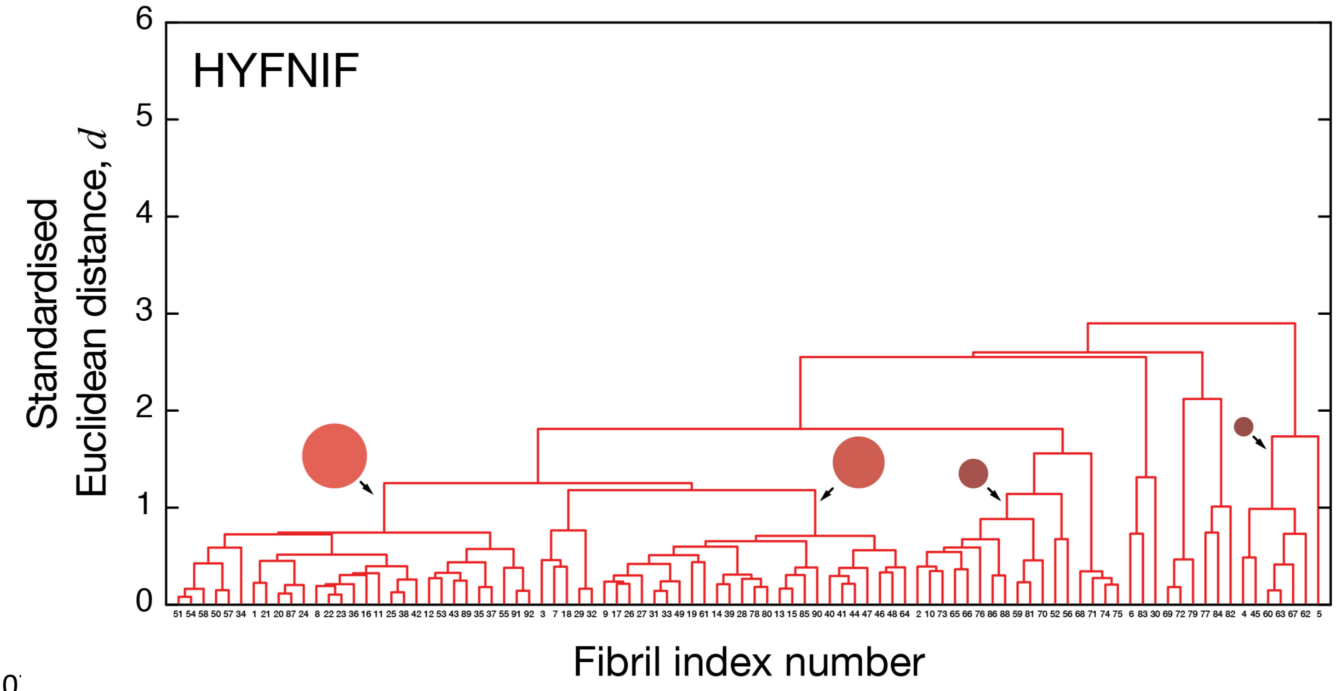
HYFNIF
Standardised Euclidean distance, $d$
0
1
2
3
4
5
6
Fibril index number

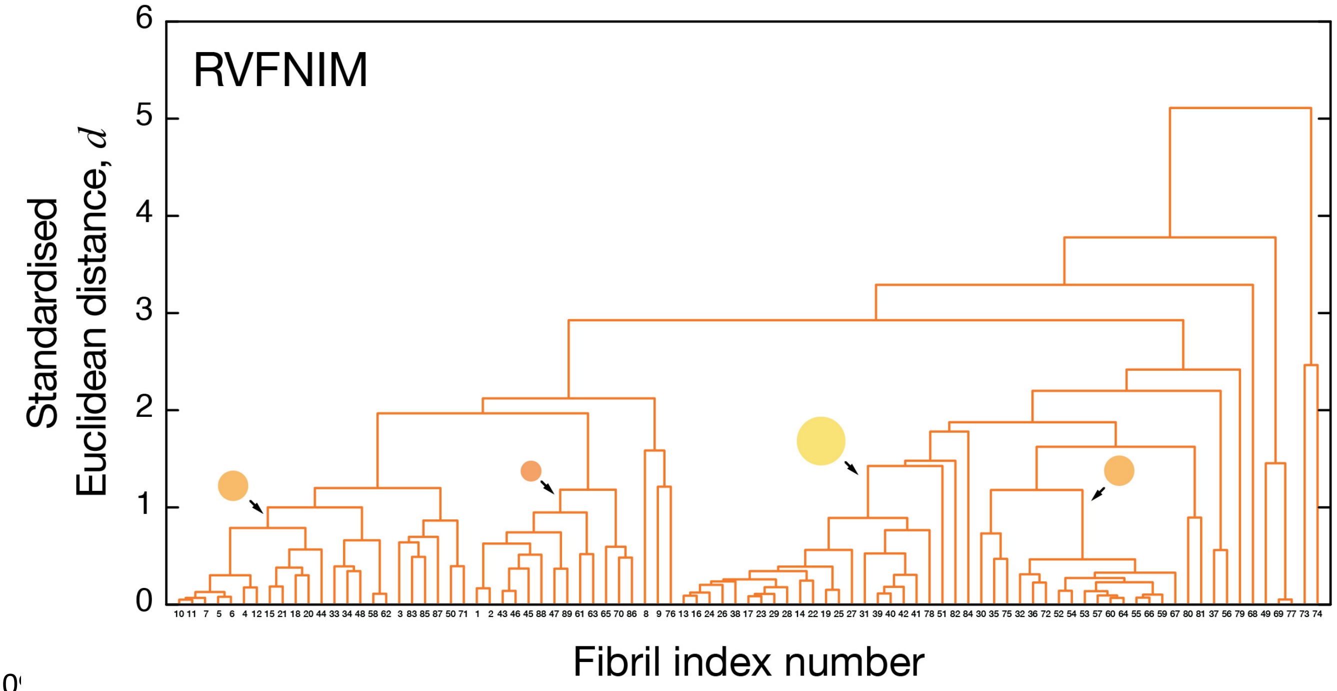
RVFNIM
Standardised Euclidean distance, $d$
0
1
2
3
4
5
6
Fibril index number

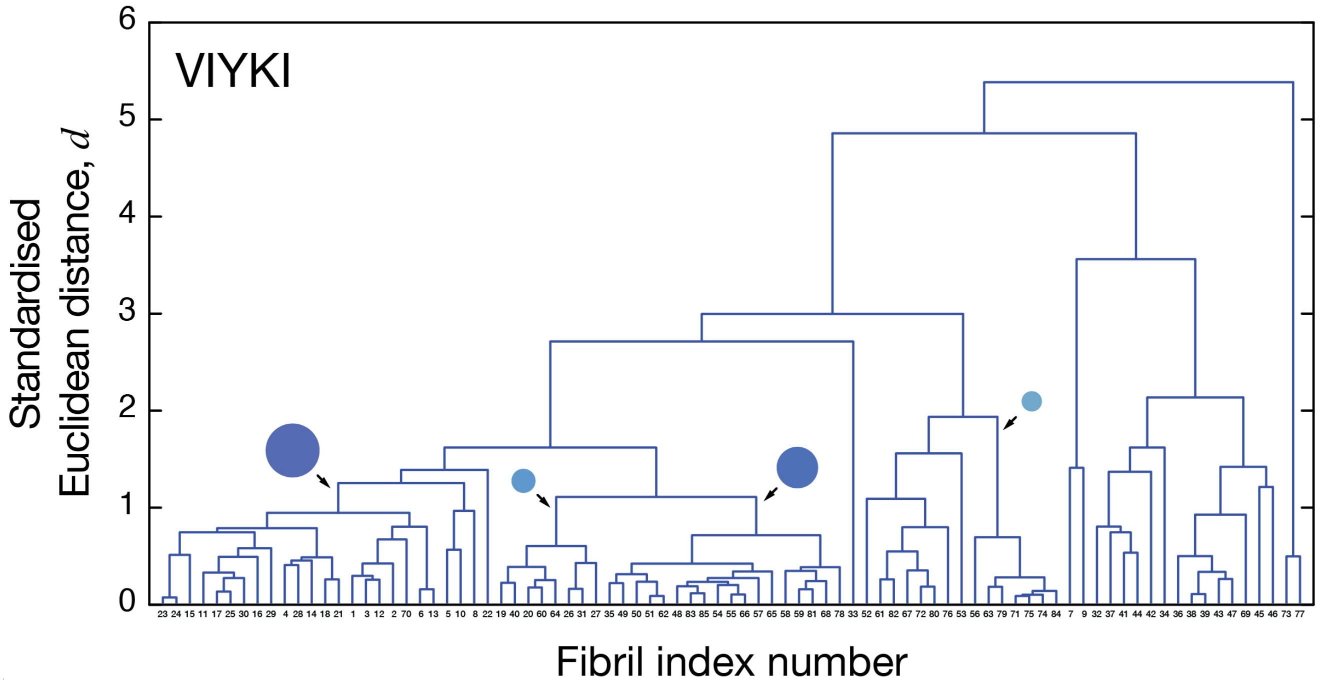

**Supplementary Figure SI 5: Analysis of structural similarity and objective classification of fibril assemblies by agglomerative hierarchical clustering.** Full dendrograms representing the full hierarchical relationship between each individual fibril is shown. For each fibril, the individual numbers shown on the x-axis are the fibril index numbers used throughout (see Supplementary Table SI 1). The x-axis represents the order in which structurally similar fibrils were grouped together, and represent the same order used in Supplementary Figures SI 1, 2 and 3. The coloured circles represent the four largest class for each sample as shown in Figure 6.

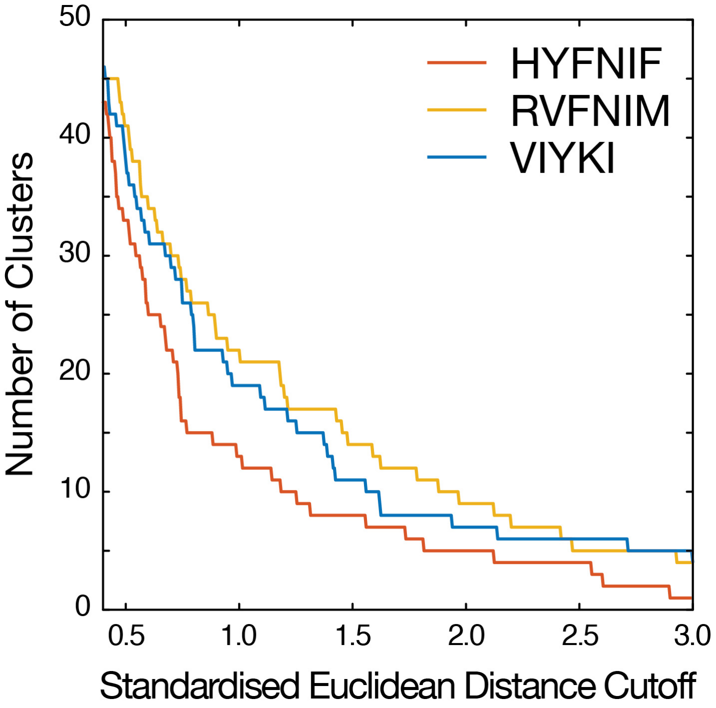

**Supplementary Figure SI 6: The number of clusters generated by agglomerative hierarchical clustering as function of standardised Euclidean distance cut-off used.** A standardised Euclidean distance cut-off of 1.0 was used to classify the fibrils in each data set (Figure 6). HYFNIF fibrils show greatest similarity as measured by the standard deviation of standardised Euclidean distances, and the data set can be described by smallest number of clusters compared to RVFNIM and VIYKI fibrils, as expected. Despite the standard deviation of standardised Euclidean distance for VIYKI fibrils is greatest of the three samples, roughly the same number of clusters if not more are required to describe the entire RVFNIM data set. This suggest that despite the VIYKI data set showing an overall greater variation, the RVFNIM fibril structures is spread across smaller distances but more evenly compared to VIYKI fibrils.
